# A compendium of co-regulated protein complexes in breast cancer reveals collateral loss events

**DOI:** 10.1101/155333

**Authors:** Colm J. Ryan, Susan Kennedy, Ilirjana Bajrami, David Matallanas, Christopher J. Lord

## Abstract

Protein complexes are responsible for the bulk of activities within the cell, but how their behavior and composition varies across tumors remains poorly understood. By combining proteomic profiles of breast tumors with a large-scale protein-protein interaction network, we have identified a set of 258 high-confidence protein complexes whose subunits have highly correlated protein abundance across tumor samples. We used this set to identify complexes that are reproducibly under- or over-expressed in specific breast cancer subtypes. We found that mutation or deletion of one subunit of a complex was often associated with a collateral reduction in protein expression of additional complex members. This collateral loss phenomenon was evident from proteomic, but not transcriptomic, profiles suggesting post-transcriptional control. Mutation of the tumor suppressor E-cadherin (*CDH1)* was associated with a collateral loss of members of the adherens junction complex, an effect we validated using an engineered model of E-cadherin loss.

## Introduction

Multi-subunit protein complexes are responsible for the bulk of the functionality of the cell (Alberts, 1998; Hartwell et al., 1999). Despite their importance to cellular function, relatively little is known about how the functionality and composition of protein complexes is altered in different cancer subtypes or in individual cancer patients. Recent examples in breast cancer suggest that even ‘housekeeping’ complexes traditionally thought of as constitutively active and essential in all cell types, such as the ribosome and the spliceosome, may become differentially expressed or differentially essential in specific contexts (Hsu et al., 2015; Pozniak et al., 2016). Consequently there is a great need to characterise the altered behavior of protein complexes in cancer.

Largely for technical and economic reasons, the large-scale molecular profiling of tumors performed over the past decade has focused on characterising changes at the genomic and transcriptomic level. Transcriptomic measurements are often used as a proxy measurement for protein expression, but most genes display only a moderate correlation between their mRNA and protein expression levels (Liu et al., 2016; Vogel and Marcotte, 2012), e.g. an average correlation of ~0.4 between mRNA and protein abundance was reported in two recent large-scale studies (Mertins et al., 2016; Zhang et al., 2016). Moreover, this correlation varies considerably between genes, with members of large protein complexes such as the ribosome and spliceosome reported to have significantly lower mRNA-protein correlation than average (Mertins et al., 2016; Zhang et al., 2016). Taken together, these observations suggest that efforts to understand altered protein complex functionality must rely on more direct measurements of protein expression. Reverse-phase protein array (RPPA) analyses have been used to quantify the expression of proteins and phosphoproteins in thousands of tumor samples, but typically these analyses only assess the abundance of ~200 (phospho)proteins and primarily focus on proteins involved in specific signaling networks (Akbani et al., 2014), limiting their wider utility for understanding protein complex regulation. Recently, advances in mass-spectrometry have enabled the quantification of thousands of proteins across large numbers of samples (Mertins et al., 2016; Pozniak et al., 2016; Riley et al., 2016; Tyanova et al., 2016; Zhang et al., 2014). These datasets permit, for the first time, a large-scale assessment of the behavior of protein complexes across different tumor samples and between different tumor types.

Here, we explore the behavior of protein complexes across 77 breast tumor proteomes (Mertins et al., 2016). We find that proteins belonging to the same complex tend to display correlated protein expression profiles across tumor samples, significantly more so than can be observed using mRNA expression profiles. We exploit this phenomenon by integrating a large-scale protein-protein interaction network with proteomic profiles of tumor samples to identify a set of protein complexes that are coherently expressed across breast tumor proteomes (BrCa-Core). These include examples of well-characterised complexes as well as those that are less well studied. As compared to a literature-curated set of protein complexes, the BrCa-Core complexes display higher correlation across additional proteomic and functional datasets. We find a number of instances where these complexes are reproducibly over- or under-expressed in specific breast cancer subtypes defined by hormone receptor status.

In assessing the BrCa-Core complexes, we find multiple instances where loss of one protein complex subunit, via gene mutation or homozygous deletion, is associated with reduced expression of additional complex subunits. This collateral subunit loss phenomenon suggests that recurrent mutations or deletions impact not only individual proteins but also the complex that they belong to. In all cases identified, this reduction of expression is evident at the protein but not mRNA level, suggesting post-transcriptional mechanisms are responsible. A notable example of this phenomenon involves E-Cadherin (*CDH1*), a tumor suppressor recurrently mutated in breast cancer (Berx et al., 1995). We find that loss of E-cadherin is associated with a reduction in protein expression of multiple members of an adherens junction complex to which it belongs. In *CDH1* mutant tumor samples, the overall expression of E-cadherin associated complex members is reduced. We replicate this finding in a *CDH1* mutant isogenic cell line, which confirms that E-cadherin loss plays a causative and not just correlative role in this reduction of expression.

## Results

### Similarity of co-expression profiles is highly predictive of protein complex membership

We first wished to assess whether known protein complexes are coherently regulated across tumor proteomes. Using the CORUM manually curated set of human protein complexes (Ruepp et al., 2010) and 77 protein expression profiles from the Cancer Genome Atlas (TCGA) breast cancer proteomics project (Mertins et al., 2016) we found that the average correlation between protein pairs in the same complex was significantly higher (Pearson’s r = 0.23) than that between random protein pairs (Pearson’s r = 0.03). We used a receiver operating characteristic (ROC) curve to assess the relationship between the similarity of protein-expression profiles and the likelihood of two proteins belonging to the same protein complex (Figure 1A). In comparison to the correlation observed using mRNA expression profiles, protein expression profiles were significantly more predictive of co-complex membership (Figure 1A) (Area Under the ROC Curve (AUC) 0.70 vs AUC 0.61). This observation is consistent with recent work that found, using tumor profiles, that protein co-expression was more predictive of general functional similarity than mRNA co-expression (Wang et al., 2017). Although co-expression calculated over the same number of samples suggested a significant advantage of proteomic profiles over mRNA profiles, the number of existing tumor with mRNA profiles far exceeds the number with proteomic profiles. Consequently it was important to determine whether the relatively small number of existing breast tumor proteomes (n = 77) could be used to predict co-complex membership with greater accuracy than the full set of TCGA tumor samples with mRNA profiles available (n=1,100)(Ciriello et al., 2015). We found that even with ~14 times as many mRNA profiles as proteomic profiles the proteomic profiles still outperformed mRNA in predicting co-complex membership (AUC 0.70 vs 0.64, Figure 1). This suggested that post-transcriptional processes such as translation and protein turnover may significantly contribute to maintaining the stoichiometry of protein complexes.

**Figure 1.**
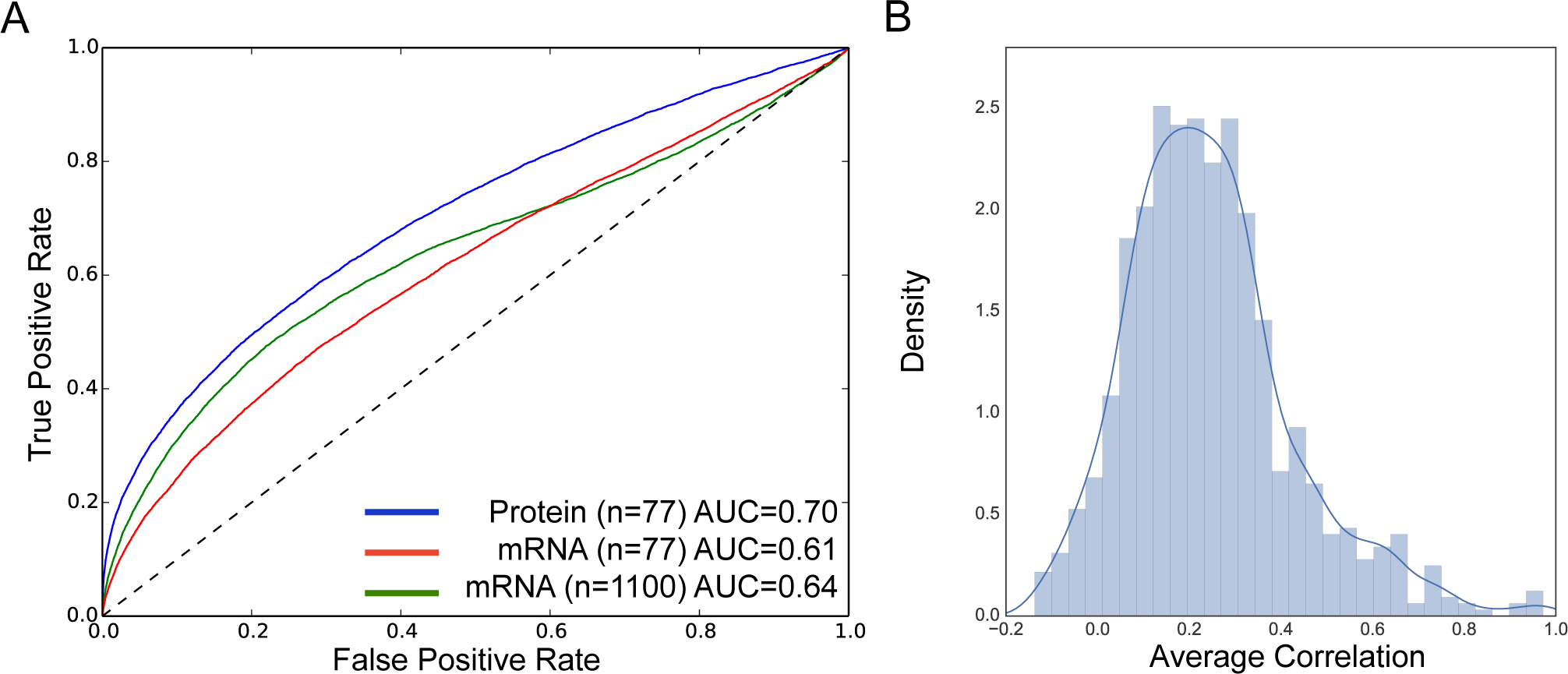
Protein co-expression is predictive of co-complex membership. A) ROC curve showing the ability of co-expression to predict co-complex membership. Blue line shows ROC curve derived from correlation calculated using protein expression profiles from 77 tumors, while the red line indicates that derived from mRNA expression profiles for the same samples. The green line indicates the co-expression correlation calculated using a larger set of 1,100 mRNA expression profiles. B) Histogram showing the average protein expression correlation within CORUM complexes. Only the average correlations from complexes with at least three members are shown in the plot.

While in general the expression of different subunits within the same CORUM complex was highly correlated, this was not the case for all complexes examined, suggesting that not all complexes are coherently regulated to a similar degree in breast cancer (Figure 1B). For example the MCM complex (Ishimi et al., 1996) had an average Pearson’s correlation of 0.96 between the protein expression profiles of its subunits, while the GAIT complex (Sampath et al., 2004) had an average correlation between subunits of -0.04. Moreover, visual exploration of the expression data suggested that there were highly-correlated groups of proteins corresponding to known complexes that were absent from the CORUM curated set. With these issues in mind, for further analysis we elected to use a data-driven approach to identify protein complexes coherently regulated in breast cancer (Figure 2).

**Figure 2.**
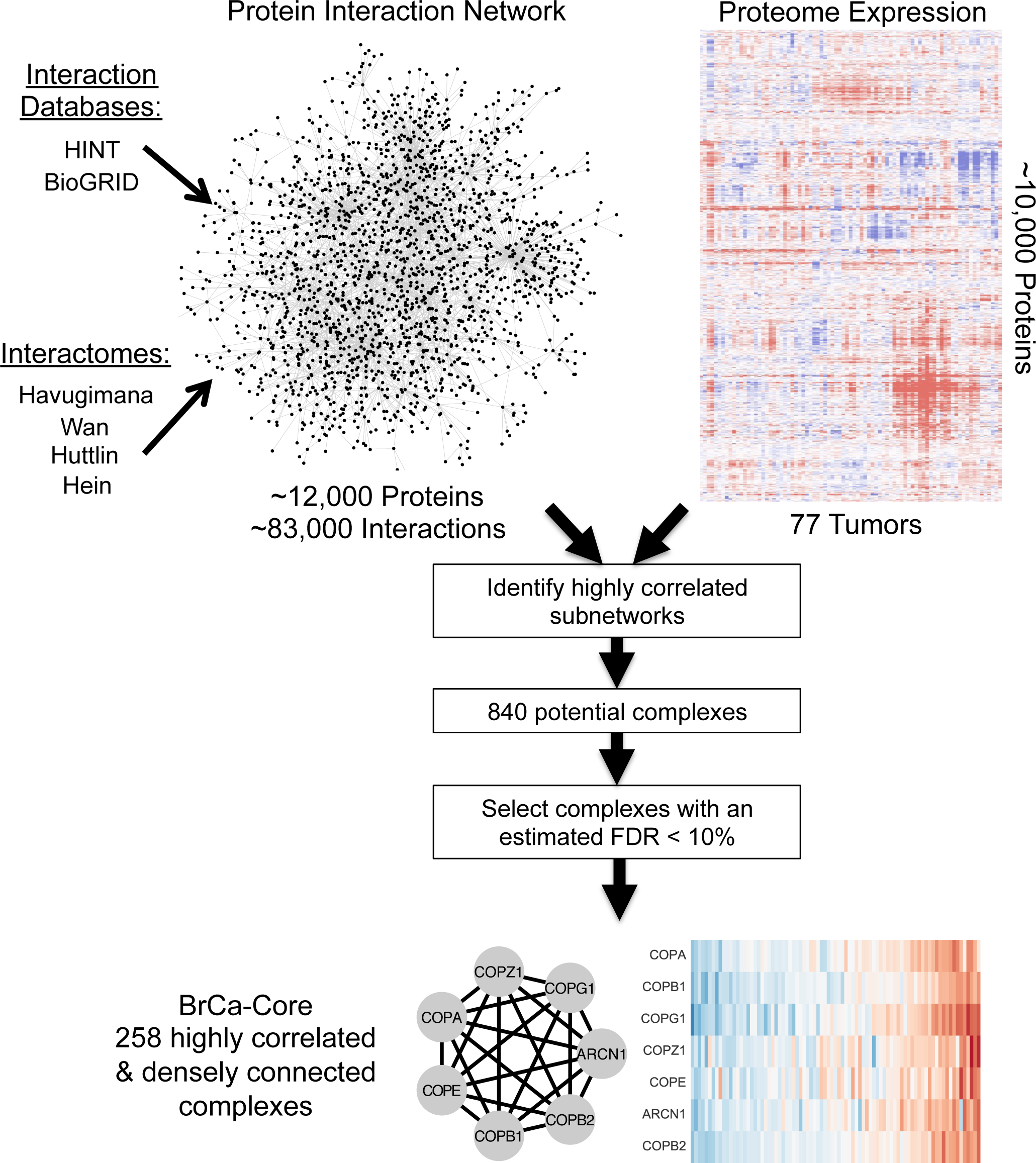
BrCa-Core complex discovery schematic. An integrated protein-protein interaction network is combined with tumor proteomic profiles to identify sets of densely connected proteins that display correlated expression profiles across tumor proteomes. By comparing the results to those derived from randomly relabeled protein interaction networks we can estimate the false-discovery rate. The BrCa-Core set contains 258 complexes at an estimated FDR of 10%.

### A compendium of protein complexes co-regulated in breast tumors

We hypothesized that by integrating large-scale protein-protein interaction networks with proteomic profiling we could identify protein complexes coherently regulated in breast tumors. We first constructed a large network of protein-protein interactions by integrating literature curated interaction databases (Chatr-Aryamontri et al., 2016; Das and Yu, 2012) with recently generated large scale high-throughput protein interaction maps (Havugimana et al., 2012; Hein et al., 2015; Huttlin et al., 2015; Wan et al., 2015). This approach generated an integrated network containing ~83,000 interactions between ~12,000 individual proteins (Figure 2).

To identify sets of genes that are densely connected on this network and display highly correlated expression profiles across multiple tumor samples we developed a machine learning approach that integrated the protein-protein interaction network with proteomic expression profiles from 77 breast tumors (Mertins et al., 2016) (Figure 2, methods). Using this approach we identified a high-confidence set of 258 complexes encompassing 1,059 distinct proteins (Figure S1, Table S1). We refer to this set of complexes throughout as BrCa-Core 1-258. The identified complexes range in size from 2 subunits to 43 subunits (mean size = 4.1) with the largest complex corresponding to the cytosolic ribosome (BrCa-Core 1). Just over half of the BrCa-Core complexes (n=131) significantly overlap with literature curated complexes annotated in CORUM (adjusted p-value < 0.05), including the COP9 signalosome (Figure 3A, BrCa-Core 17) (Seeger et al., 1998) and the conserved oligomeric Golgi (COG) complex (Figure 3B, BrCa-Core 14) (Ungar et al., 2002). Some of the BrCa-Core complexes encapsulated protein complexes already annotated in CORUM along with additional subunits – for example BrCa-Core 47 included the CORUM annotated origin-recognition 2-5 complex (ORC 2-5) (Dhar and Dutta, 2000) with the addition of LWRD1 (also known as ORCA); ORCA interacts with the ORC complex and stabilizes binding of the complex to chromatin (Shen et al., 2010) (Figure 3C). Complexes identified in BrCa-Core but absent from the CORUM human complex set include the COPI-vesicle coat complex (Figure 3D, BrCa-Core 25), a variant of the endosome-associated recycling protein (EARP) complex that includes all four EARP subunits along with the more recently identified EARP interactor TSSC1 (Gershlick et al., 2016; Schindler et al., 2015)(Figure 3E, BrCa-Core 48), and a complex containing the majority of subunits of the newly identified ‘Commander’ (COMMD/CCDC22) complex (Figure 3F, BrCa-Core 26) (Starokadomskyy et al., 2013) not included in CORUM but recently shown to be highly conserved across metazoans (Wan et al., 2015).

**Figure 3.**
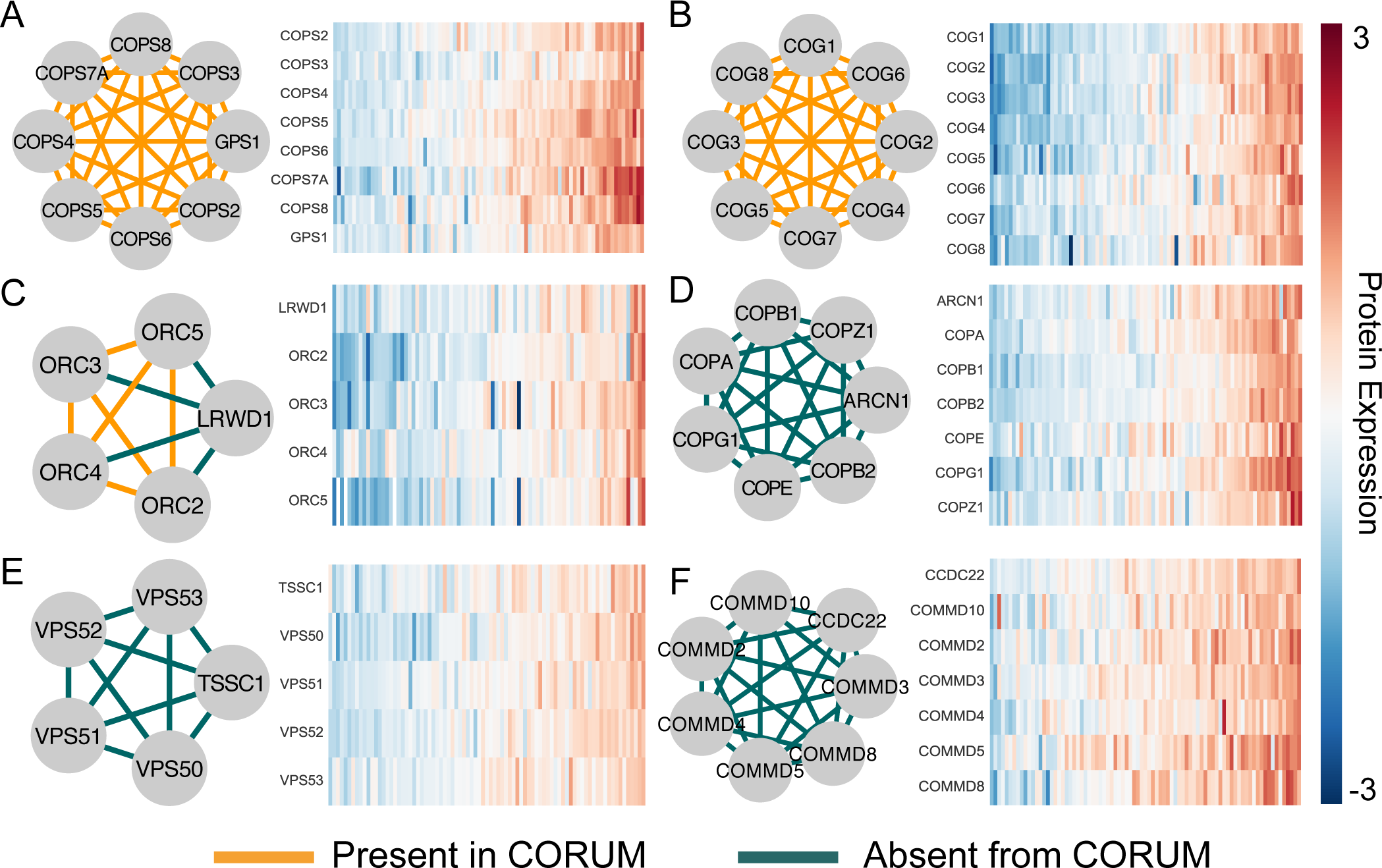
Complexes in BrCa-Core. A) BrCa-Core 17 - the COP9 signalosome. All cocomplex relationships are also found in the equivalent CORUM complex. The heatmap in the right shows protein expression of all subunits across 77 breast tumour proteomes. These have been sorted based on the mean abundance of all subunits B) BrCa-Core 14 - the conserved oligomeric golgi (COG) complex. All cocomplex relationships shown are also found in the equivalent CORUM complex. C) BrCa-Core 47 - contains ORC2-5 complex found in CORUM with the addition of LRWD1 D) BrCa-Core 25 - the COPI complex E) BrCa-Core 48 - the EARP complex with the recently identified EARP interactor TSSC1 F) BrCa-Core 26 - the Commander complex

The majority of BrCa-Core complexes have significant overlap with specific Gene Ontology Cellular Component and Biological Process terms, suggesting common localization and functionality respectively (196 complexes enriched in GO-CC terms, 218 enriched in GO-BP terms, both at adjusted p-value < 0.05) (Table S1). Like known protein complexes, pairs of proteins assigned to the same BrCa-Core complex were significantly more likely than random protein pairs to be frequently mentioned together in the literature (Odds-Ratio 176, p < 1x10^-16^, Fisher’s exact test) and to display similar patterns of conservation across species (Odds-Ratio 280, p < 1x10^-16^, Fisher’s exact test). When pairs of proteins annotated to the same CORUM complex were excluded from this analysis, i.e. when analyzing only new co-complex pairs identified within BrCa-Core set, we still observed a significant enrichment for protein pairs to be frequently mentioned together in the literature (Odds-Ratio 95, p < 1x10^-16^, Fisher’s exact test) and to display similar patterns of conservation across species (Odds-Ratio 335, p < 1x10^-16^, Fisher’s exact test). This suggests that the newly identified complexes display similar functional cohesion to those in the CORUM database.

As our method exploited the correlation between protein expression profiles to identify complexes, we expected the average correlation across the TCGA BrCa proteomes within the BrCa-Core complexes to be significantly higher than random protein pairs. This was indeed the case with an average correlation of 0.62 compared to an expected correlation of 0.03. This correlation was significantly higher than the average of pairs in our integrated protein interaction network (0.12) or pairs within CORUM complexes (0.23). To assess whether the same higher correlation could be observed in additional tumor proteomic resources we analyzed two additional breast tumor proteomic datasets (Pozniak et al., 2016; Tyanova et al., 2016). Tyanova *et al* contains proteomic profiles of 40 tumor samples from diverse breast cancer subtypes, while Pozniak *et al* contains proteomes of 66 primary luminal-subtype breast tumors or metastatic lesions. Again the average within BrCa-Core correlation was significantly higher in these datasets than the correlation observed between CORUM complex pairs or among random protein-protein interaction pairs (Table 1).

**Table 1.** Correlation of BrCa-Core co-complexed pairs in additional resources. Shown is the average correlation between pairs of proteins annotated to the same protein complex (CORUM or BrCa-Core) or identified as interacting in the integrated protein-protein interaction network (PPI Pairs). Correlation is calculated across three breast tumour proteomic datasets (TCGA, Tyanova, Pozniak) and an shRNA screen of breast tumour cell lines (Marcotte).

|  | **TCGA**<br>**Proteomics** | **Tyanova**<br>**Proteomics** | **Pozniak**<br>**Proteomics** | **Marcotte**<br>**shRNA** |
| --- | --- | --- | --- | --- |
| **PPI Pairs** | 0.12 | 0.12 | 0.10 | 0.06 |
| **CORUM** | 0.20 | 0.19 | 0.14 | 0.08 |
| **BrCa-Core** | **0.62** | **0.32** | **0.28** | **0.25** |

The tendency of pairs of proteins within the same complex to display similar phenotypes when inhibited has been well established in the literature, especially in model organisms such as budding yeast and *Caenorhabditis elegans* (Sharan et al., 2007; Wang and Marcotte, 2010). This phenomenon has been used to predict novel phenotypes for members of protein complexes via the ‘guilt-by-association’ principle (Sharan et al., 2007; Wang and Marcotte, 2010). To assess whether the BrCa-Core complexes also displayed a similar tendency, we analyzed the results of a recently published large-scale shRNA screen in 77 breast tumor cell lines (Marcotte et al., 2016). This screen measured the cell growth inhibition effects of shRNAs targeting ~16,000 genes in each cell line. We expected that shRNAs targeting members of the same complex would display correlated essentiality profiles (i.e. would inhibit tumor cell lines in a similar fashion) and we found that this is indeed the case. Pairs of proteins in the same BrCa-Core complex displayed a significantly higher correlation (~0.25) than random pairs (~0.0) and than pairs annotated to the same CORUM complex (~0.14), suggesting that they are more phenotypically similar (Table 1).

It is reasonable to ask why the majority of CORUM protein complexes were not more highly co-expressed across samples. If all members of the complex function as a single unit, why would they not display highly correlated protein expression? One explanation is that protein complexes exist in multiple isoforms, with the exact composition varying across cell types and conditions. Consequently pairs of proteins within the same complex may display lower than expected correlation due to one of them functioning in additional protein complexes. Indeed in the CORUM the average protein belongs to 3 distinct complexes, while by design in BrCa-Core each protein was assigned to a single complex based on highly correlated expression with other members. Previous work in yeast has suggested that the subunits of protein complexes can be divided into two groups - cores (proteins found in the majority of complex isoforms) and attachments (proteins found in a small number of isoforms)(Gavin et al., 2006). Some pairs of attachment proteins are often found together in multiple complexes, and these have been referred to as ‘modules’(Gavin et al., 2006). One explanation for the higher correlation seen within the BrCa-Core complexes is that they preferentially identify sets of core proteins or modules. Consistent with this we found that pairs of proteins annotated together in two or more CORUM complexes (suggesting they resemble ‘modules’) were significantly more likely to be identified together in a BrCa-Core complex (Odds-Ratio = 2.2) as were pairs always found in the same CORUM complex (consistent with them being either ‘modules’ or ‘cores’)(Odds-Ratio =1.8).

### Differential expression of protein complexes in breast cancer subtypes

At the molecular level breast cancer is a very heterogenous disease, with each tumor displaying a unique genetic and epigenetic profile. Despite this heterogeneity, molecular biomarkers can be used to classify tumors with similar molecular profiles into subtypes (Onitilo et al., 2009; Perou et al., 2000; Sorlie et al., 2001). These molecular subtypes, to some extent, display different survival outcomes and different responses to targeted therapies (Onitilo et al., 2009; Perou et al., 2000; Sorlie et al., 2001). Although a variety of subtypes have been defined using mRNA expression profiles and/or genomic classifiers, the biomarkers used most commonly in the clinic are the estrogen receptor (ER), progesterone receptor (PR) and human epidermal growth factor receptor 2 (ERBB2/HER2), often measured using immunohistochemistry (IHC) (Onitilo et al., 2009). The status of these markers can be used to classify breast tumors into four broad subtypes – triple negative (TNBC, ER^-^/PR^-^/HER2^-^), HER2 positive (ER^-^/PR^-^/HER2^+^), ER^+^/PR^+^/HER2^-^ and ER^+^/PR^+^/HER2^+^ (Onitilo et al., 2009). Respectively, these subtypes somewhat overlap with Basal, HER2^+^, Luminal A and Luminal B “intrinsic” subtypes defined using mRNA expression profiles (Onitilo et al., 2009; Perou et al., 2000; Sorlie et al., 2001). To better understand how breast cancer subtypes might influence protein complexes (and *vice-versa*) we assessed the relationship between BrCa-Core protein complex expression and IHC-defined subtypes. To enable the identification of reproducible associations between subtypes and protein complex abundance we focused on those subtypes with reasonable representation in both the TCGA Mertins *et al* (Mertins et al., 2016) resource and the dataset of Tyanova *et al*(Tyanova et al., 2016). As Tyanova *et al* has only 2 samples that are ER^+^/PR^+^/HER2^+^ we excluded samples of this subtype from our analysis, and focussed on three subtypes common to both resources – HER2+ (ER^-^/PR^-^/HER2^+^), ER^+^ (ER^+^/PR^+^/HER2^-^) and triple negative (ER^-^/PR^-^/HER2^-^).

Using the TCGA dataset and the BrCa-Core complexes, we discovered 82 associations between subtype and complex abundance at a false discovery rate (FDR) of 10% (Table S2, Figure S2). At the same FDR threshold we found 7 associations using the CORUM complex set, highlighting the advantage of using BrCa-Core for this analysis. Due to differences in coverage of protein complex subunits, not all of the 82 associations could be tested in the Tyanova *et al* dataset (Tyanova et al., 2016).

Of the 59 associations that could be tested 27 were observed in the Tyanova *et al* dataset at the same FDR of 10% (Table S2). In general the effect sizes and directions across the two datasets were highly correlated (Spearman’s r = 0.69, p < 1x10^-8^) suggesting that with larger sample sizes additional associations between subtype and complex abundance could be replicated. Examples of differentially expressed complexes replicated in the Tyanova *et al* dataset are presented in Figure 4. Triple-negative breast tumors were associated with increased expression of a number of complexes involved in DNA replication including the replication factor C complex (BrCa-Core 21) and the MCM complex (BrCa-Core 28) (Figure 4). Different members of the MCM complex (MCM2 and MCM4) have previously been identified as markers of proliferation, associated with poorer survival outcomes in breast cancer and shown to have higher expression in ER negative breast tumors (Joshi et al., 2015; Kwok et al., 2015). ER^+^ tumors were associated with decreased expression of two complexes involved in antigen processing (BrCa-Core 59 and 193) consistent with data suggesting that expression of antigen presentation human leukocyte antigen (HLA) molecules is lower in the ER^+^ subtype (Chung et al., 2017; Lee et al., 2016). HER2^+^ tumors were associated with increased expression of two complexes involved in Golgi transport associated vesicle coating (BrCa-Core 25 and 42). It is not immediately obvious why *HER2* amplification would be associated with an increased expression of complexes involved in vesicle transport, but the association is evident across both patient cohorts (Figure 4, Figure S2, Table S2).

**Figure 4.**
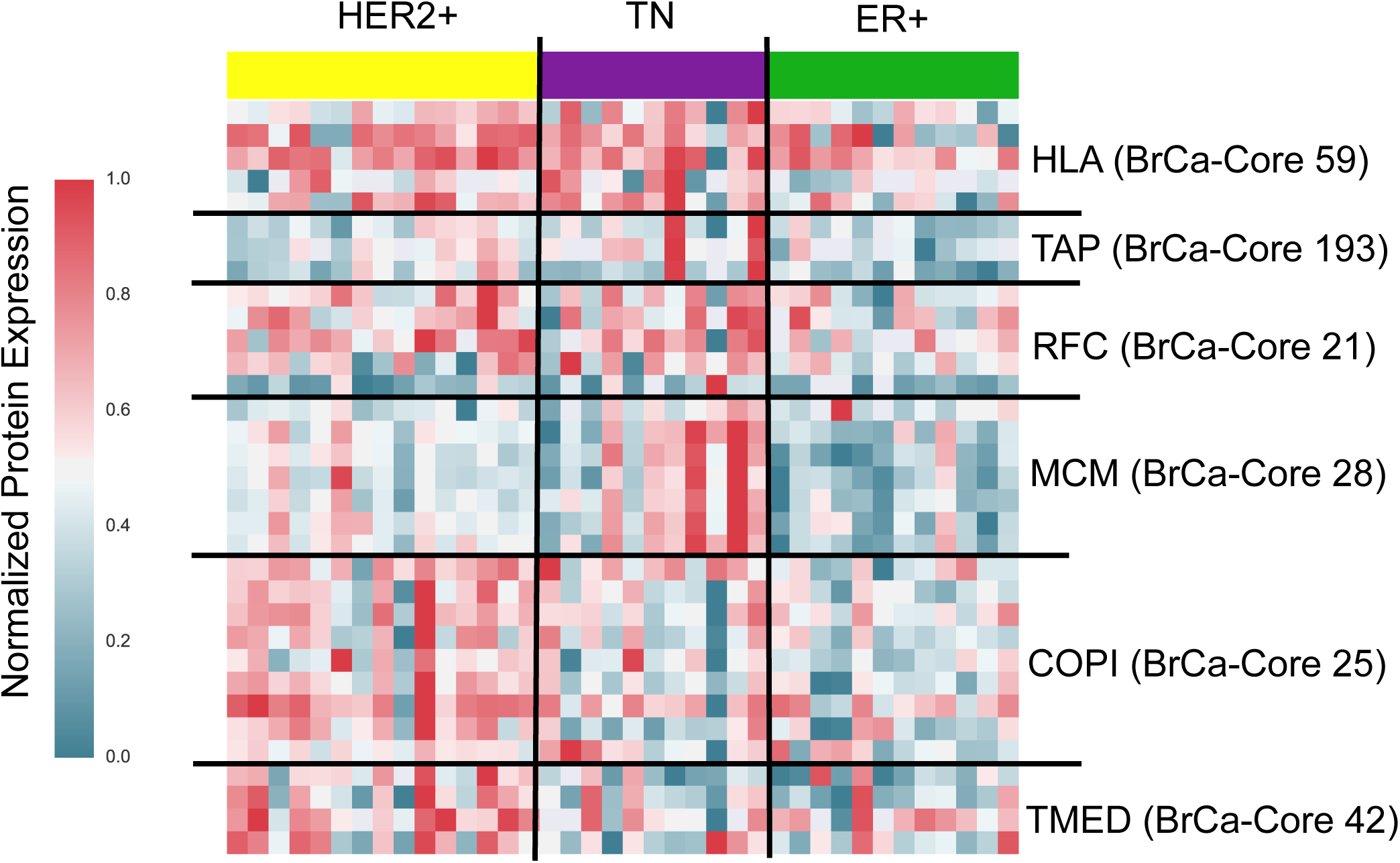
Subtype specific complex expression. Heatmap displaying protein expression levels of specific BrCa-Core complexes. Tumor samples are grouped according to subtype (using IHC markers), indicated on top of the heatmap. Genes are grouped into specific complexes indicated on the right of the heatmap. Shown are the expression levels taken from Tyanova *et al* (used for validation). These expression levels have been normalized such that the maximum expression level is 1 and minimum is 0. Heatmap for the discovery dataset (Mertens *et al*) is shown in Figure S2.

### The impact of subunit loss on protein complex expression

An implication of highly correlated protein expression within a protein complex is that loss of protein expression of one subunit might frequently be associated with reduced protein expression of other complex subunits (Figure 5A). Such a reduction in expression may occur through reduced transcription, reduced translation, or an increase in protein degradation. Consequently genetic events that reduce protein expression of one subunit, such as mutation or homozygous deletion, may be associated with a collateral reduction (in *trans*) of protein expression of other subunits or indeed the entire complex (Figure 5A). To test whether this is the case we identified five genes that are members of BrCa-Core complexes whose mutation or deletion is associated with a nominally significant (p < 0.05, Mann Whitney U test) reduction in expression of their encoded proteins (*CDH1 (E-cadherin), PBRM1, CYFIP2, GLUD1, EXOC2*). We then asked whether mutation or deletion of these genes was also associated with overall reduction in protein expression of the complex that they belong to. In all five cases we found that loss of one subunit was associated with a reduction in the protein expression of additional complex subunits. For instance homozygous deletion or mutation of *EXOC2* was associated with decreased proteomic abundance of EXOC2 and an overall reduction in the protein expression of multiple members of the exocyst complex (Matern et al., 2001) to which it belongs (including EXOC3, EXOC6, EXOC7, and EXOC8) (Figure 5B, BrCa-Core 27). While loss of *EXOC2* was also associated with a reduction of EXOC2 mRNA expression, no reduction was observed for other protein complex subunits at the mRNA level (Figure S3A) suggesting that the reduction in protein expression levels is caused by post-transcriptional mechanisms. Furthermore, the correlation between complex subunits was significantly higher at the protein expression than mRNA level (Figure S4A) suggesting these post-transcriptional mechanisms may contribute to the coherent protein expression of the complex. Similarly we found that mutation/deletion of *PBRM1*, a component of the PBAF chromatin remodeling complex most frequently mutated in renal carcinoma (Varela et al., 2011), was associated with a reduction of the expression of a complex containing the additional PBAF subunits SMARCC2, ARID2 and BRD7 (Kaeser et al., 2008) (Figure 5A, BrCa-Core 60). Again this reduced expression was evident at the protein but not mRNA level (Figure S3B) and the within complex correlation was significantly higher using protein rather than mRNA expression profiles (Figure S4B). Most intriguingly we found that mutation of the tumor suppressor *CDH1* (E-cadherin) was associated with a decreased abundance of both the E-cadherin protein and additional members of an adherens junction complex to which it was assigned in BrCa-Core (BrCa-Core 30). All proteins in this complex have highly correlated protein expression with E-cadherin (average Pearson’s correlation 0.65, Figure S4C) and four of the complex subunits (E-cadherin, CTNNA1, CTNNB1, and CTNND1) have a significant (Mann-Whitney p < 0.05) decrease in expression in *CDH1* mutant samples (Figure 5D). In contrast, the average mRNA correlation of all subunits with CDH1 was low (Pearson’s correlation 0.08, Figure S4C) with one subunit (CTNNB1) displaying weakly negative correlation with E-cadherin (Pearson’s correlation -0.16, Figure S4C). None of the subunits other than E-cadherin itself display a significant relationship between *CDH1* mutation status and mRNA expression. As loss of E-cadherin is a major driver event in breast cancer, mutated in ~11% of all breast tumors and over 50% of invasive lobular breast tumors (Berx et al., 1995; Ciriello et al., 2015; Michaut et al., 2016), we focused on the consequences of *CDH1* loss for further analysis.

**Figure 5.**
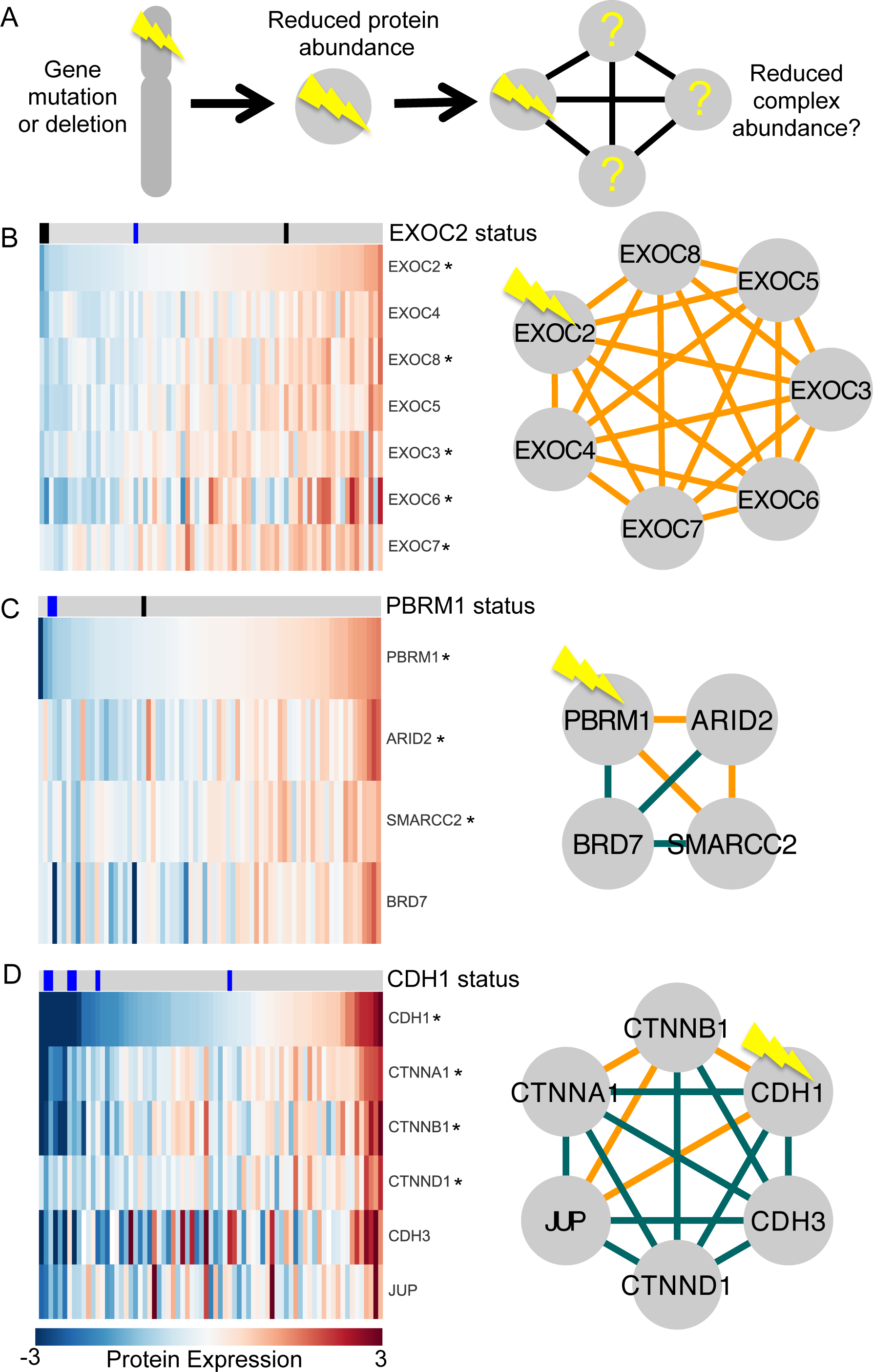
Subunit loss is associated with a reduction in protein complex expression. A) Model displaying potential series of events – mutation or deletion of one subunit is associated with reduced protein abundance of that subunit, and potentially a reduction in expression of the entire complex. B) Mutation or deletion of *EXOC2* is associated with a reduction in protein expression of the exocyst complex (BrCa-Core 27). *EXOC2* mutation (blue) or deletion (black) is indicated in the top row. Proteomic samples have been sorted according to EXOC2 abundance. Genes marked with a star indicate those whose proteomic abundance is significantly lower (one-sided Mann Whitney test, p<0.05) in samples with *EXOC2* mutation/deletion. C) Mutation or deletion of *PBRM1* is associated with a reduction in protein expression of a PBAF-like subcomplex (BrCa-Core 60). Legend as for A. C) *CDH1* mutation is associated with a reduction in protein expression of an adherens-junction complex (BrCa-Core 30). Legend as for A.

### E-cadherin loss causes reduced expression of adherens junction complex members

Our initial analysis of the 77 tumor samples with mass-spectrometry derived proteomic profiles available suggested that *CDH1* mutation is associated with a decrease in the proteomic abundance of E-cadherin itself and additional members of the adherens junction complex to which it is assigned in BrCa-Core (BrCa-Core 30). Three of the proteins in this complex (E-cadherin / CTNNA1 / CTNNB1) have also been measured using the RPPA method permitting us to assess the impact of *CDH1* mutation on protein abundance measured using an orthogonal approach. RPPA measurements were available for E-cadherin and CTNNB1 in 760 tumor samples and CTNNA1 for 64 tumor samples. We found that *CDH1* mutation was associated with a significant reduction in abundance of all three proteins (Figure 6A). In contrast, when looking at mRNA expression, we found that *CDH1* mutation was associated with reduced expression of CDH1 itself but no reduction in mRNA expression of *CTNNA1* or *CTNNB1*. This suggests that E-cadherin loss is associated with a reduction in the abundance of other complex subunits via post-transcriptional mechanisms (e.g. reduced translation or increased degradation).

One limitation of this analysis is that it identifies correlative rather than causal associations – it demonstrates that loss of one subunit is associated with reduced expression of other subunits, but it does not demonstrate a causal effect. It is of course possible that some additional factor causes reduction in expression of the entire adherens junction complex rather than the mutation of a single subunit such as *CDH1*. To establish causality we used mass spectrometry to measure differential protein expression in a pair of isogenic breast cancer cell lines (MCF7) with CRISPR-Cas9 engineered *CDH1* loss (Bajrami *et al,* submitted). *CDH1* mutations in breast tumors frequently occur in lobular breast tumors, which tend to be ER^+^, have a luminal A intrinsic subtype and often harbor *PIK3CA* mutations (Ciriello et al., 2015). We therefore inactivated *CDH1* in the MCF7 cell line, which is ER^+^, luminal A, and *PIK3CA* mutant (Bajrami *et al,* submitted). We performed label-free protein quantification of whole protein lysates in parental (*CDH1* wild type) and *CDH1* defective daughter cells. This resulted in the quantification of ~5,100 proteins (Table S3). We found 91 proteins with significantly lower protein abundance in the Ecadherin defective model (p < 0.005, FDR = ~8%) including five of the six adherens junction complex subunits (E-cadherin, CTNNA1, CTNNB1, CTNND1, JUP) (Figure 6B), suggesting that *CDH1* mutation plays a causative role in the reduction of their protein abundance. In contrast to what we observe in the tumor proteomes, in the MCF7 E-cadherin null model we observed an increase in the expression of CDH3 (Pcadherin) (Figure 6B), perhaps an example of ‘cadherin switching’ specific to this model(Cavallaro et al., 2002; Wheelock et al., 2008).

**Figure 6.**
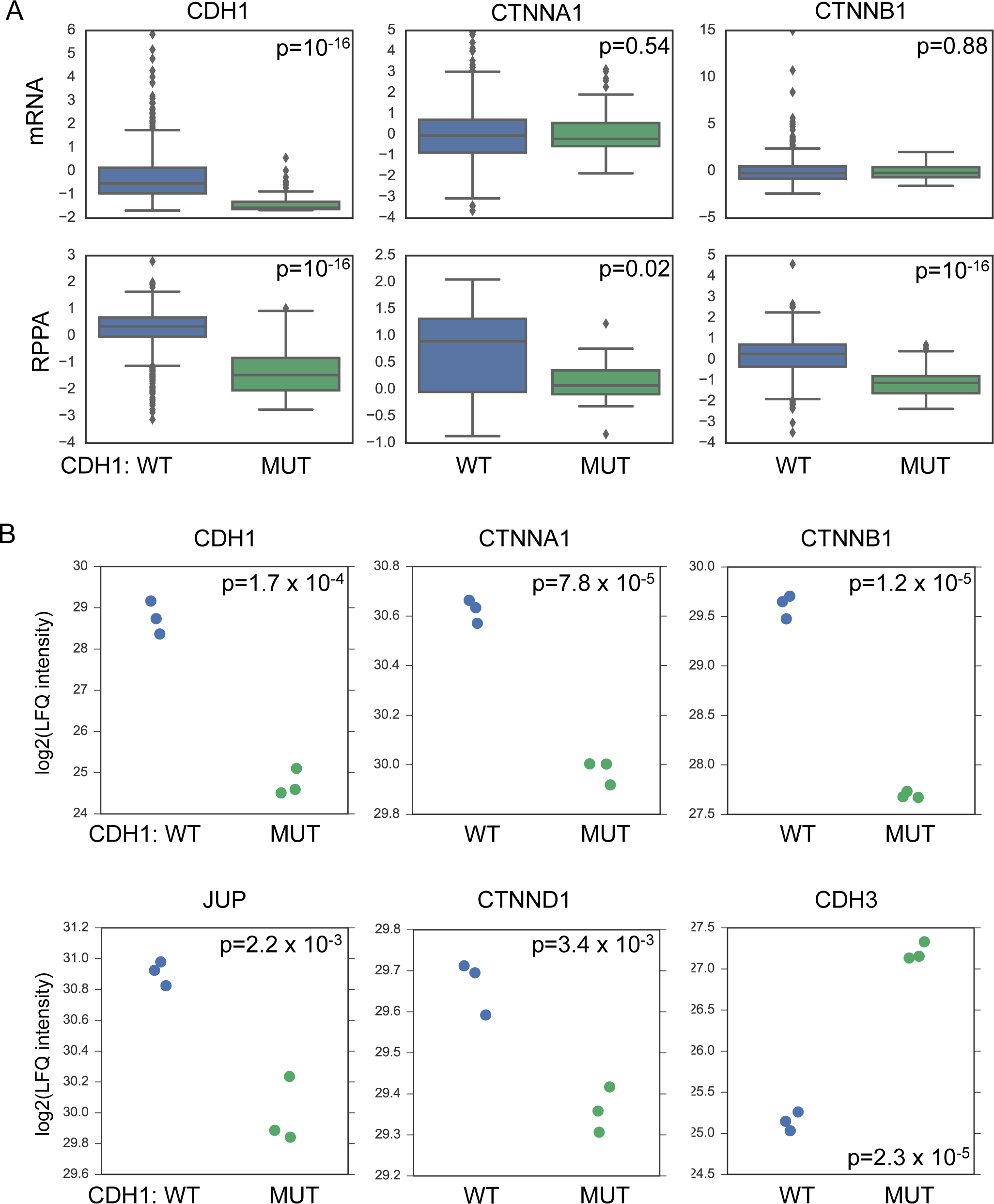
E-cadherin loss is associated with reduced expression of an adherens junction complex. A) In tumor samples *CDH1* mutation is associated with a decrease in mRNA and protein expression of CDH1, but only of protein expression for CTNNA1 and CTNNB1. All expression and RPPA measurements are Z-scores. Box plots show median and interquartile range. P-values calculated using a Mann-Whitney test. mRNA measurements for all three genes were available for 992 tumors, RPPA data for CDH1 and CTNNB1 were available for 760 tumors, while RPPA data for CTNNA1 was available for only 64 tumors. B) Protein expression measured in a pair of isogenic MCF7 cell lines that differ by *CDH1* status. Shown are the log2 Label Free Quantification intensities. P-values are calculated using a two-sided heteroscedastic t-test.

The decreased abundance of five of the six BrCa-Core adherens junction complex members in the MCF7 model was a significant enrichment over random expectation (Odds Ratio=280, p-value=10^-8^ Fishers Exact Test). To test whether our approach missed additional collateral loss events associated with *CDH1* mutation we assembled a list of 95 E-cadherin protein-protein interaction partners from CORUM (18 co-complexed subunits), BioGRID (89 protein-protein interaction partners) and HINT (15 co-complex interaction partners). Aside from the five members of the adherens junction complex in BrCa-Core, none of the known E-cadherin interaction partners displayed a significant reduction in protein abundance in the E-cadherin defective model. This suggested that our data-driven approach effectively identified the specific subunits of the adherens-junction complex whose expression is reduced by *CDH1* mutation in breast cancer.

## Discussion

Here we have used a data-driven approach to identify BrCa-Core, a set of 258 protein complexes coherently regulated across breast tumor proteomes. We found that, compared to literature curated complexes, BrCa-Core complexes are more coherently expressed in additional proteomics experiments and are also more likely to be co-essential in breast tumor cell lines. BrCa-Core includes known and potentially novel complexes, and provides additional supporting evidence for protein complexes recently identified in the literature (such as the inclusion of the TSSC1 subunit in the EARP complex)(Gershlick et al., 2016).

Using the TCGA breast tumor proteomics data (Mertins et al., 2016) we found that certain BrCa-Core complexes appear differentially expressed in particular breast tumor subtypes defined according to hormone receptor and HER2 status. We were able to reproduce a number of these associations in an additional breast tumor proteomic dataset (Tyanova et al., 2016). These two datasets were generated using different experimental platforms and different computational analysis pipelines. Furthermore they contain different proportions of the different molecular subtypes, and consequently the reproduction of associations was far from guaranteed. Nonetheless, our initial analysis suggests that a significant fraction of the subtype-specific complex associations observed in one cohort can also be replicated in the other.

We did not assess whether the expression of specific BrCa-Core complexes could be used to stratify patients to predict survival outcomes or response to specific therapies. Whilst this may be a possibility, the total number of patients with proteomic profiling available is still relatively small and precludes meaningful survival analysis. For example, eight of the 77 patients profiled in the TCGA Mertins *et al* resource had died at the time of publication (Mertins et al., 2016), a number which does not provide sufficient statistical power to establish relationships between protein expression and clinical outcome.

We found that in general, correlation between protein expression profiles predicts co-complex membership better than correlation between mRNA expression profiles. One factor that contributes to this improved correlation is the collateral loss phenomenon we observe - when one subunit of a complex is lost via deletion or mutation, a collateral loss in the protein expression of additional complex members is observed. This collateral loss is not observed at the mRNA level, and consequently complexes that experience collateral loss display higher correlation at the protein than mRNA level. There are likely many other factors that contribute to maintaining the coherent expression of protein complexes across tumors, including dosage compensation of copy number amplified genes, a phenomenon that has been observed in cancer cell lines and in models of aneuploidy in yeast (Dephoure et al., 2014; Geiger et al., 2010; Ishikawa et al., 2017; Stingele et al., 2012).

We have focused primarily on how deletions or mutations of protein complex subunits may impact the abundance of their associated complexes. Recently, by analyzing protein quantitative trait loci (pQTLs) in outbred mice, Chick *et al* (Chick et al., 2016) identified genetic variants that alter the abundance of a protein in *cis* and its interaction partners in *trans*. A notable example of this involved the CCT complex – a non-coding variant in the promoter region of *CCT6A* was associated with a reduction in the expression of the CCT6A protein and additional members of the CCT complex. This suggests that genetic alterations more subtle than the mutations and deletions analysed here may also cause collateral loss effects on protein complex members. A number of tumor suppressors, including *CDH1*, are subject to recurrent hypermethylation and therefore it would be worth testing if this mechanism of gene silencing can result in the collateral loss of complex subunits. Visual inspection of Figure 5D suggests that samples with reduced E-cadherin protein expression, even in the absence of CDH1 mutation, have reduced expression of the entire adherens junction complex. This suggests that alternative gene silencing effects may also cause collateral loss events.

We have not addressed here the mechanisms responsible for the collateral loss phenomenon, although the observation that the reduction in protein expression levels is not evident at the mRNA level suggests posttranscriptional mechanisms must be responsible. Perhaps the simplest explanation is that loss of one subunit prevents a complex from assembling, and consequently there is an increase in the proteasomal degradation of unbound subunits. However, the specific example of the E-cadheren containing adherens junction complex suggests that the story may be more complicated. In the MCF7 E-cadherin null model we observe a decrease in the abundance of most adherens junction members, but an increase in abundance of the CDH1 paralog CDH3. CDH1 (E-cadherin) and CDH3 (P-cadherin) are both members of the cadherin family, and this is potentially a case of ‘cadherin switching’ – where loss of one cadherin is associated with a compensatory increase in the expression of an alternative cadherin(Cavallaro et al., 2002; Wheelock et al., 2008). The reduction in expression of other members of the adherens junction complex suggests that if cadherin switching is occurring in the MCF7 line it does not completely compensate for E-cadherin loss and perhaps results in a less stable adherens junction complex. Although cadherin switching between E-cadherin and P-cadherin has been observed in breast tumors previously (Palacios et al., 1995), recent work suggests that in lobular breast tumors (where CDH1 mutation is most common) expression of Pcadherin is a rarity (Turashvili et al., 2011). It is possible that alternative cadherin switching (such as between E-cadherin and N-cadherin) is more common in breast tumors but we have not addressed that here due to the absence of N-cadherin from the tumor proteomics dataset.

Generally we have focused on the behavior of coherently expressed protein complexes across breast tumor samples. This approach has a number of advantages - in particular it allows us to see how different complexes behave as a single unit within molecularly defined groups of tumors. We can identify instances where an entire protein complex is up or down regulated in the presence of specific molecular markers. A disadvantage of this approach is that we cannot identify when different variants / isoforms of a protein complex become more or less abundant in specific conditions. We have overlooked such events here, but recent work in cancer cell lines and mouse fibroblasts suggest that they may be relatively common and merit further investigation (Ori et al., 2016).

We expect that the BrCa-Core complexes will be useful for the analysis of additional proteomic and functional datasets and make the full list of complexes available in Table S1. We also anticipate that the complex identification approach described here will be useful for the analysis of other large-scale proteomic datasets, such as those from other tumor or cell line profiling projects (Lawrence et al., 2015; Zhang et al., 2014; Zhang et al., 2016), and we make our code available to facilitate such efforts.

## Materials and Methods

Identifiers in all protein-protein interaction networks, protein expression datasets, and validation sets were converted to ENTREZ gene IDs. In cases where a particular gene or protein could not be matched to an ENTREZ gene ID it was discarded from further analyses.

### Protein Expression Datasets

For the primary analysis we used the breast tumor proteomics dataset from the TCGA CPTAC project (Mertins et al., 2016). Only samples that passed the authors’ quality control (77 samples, 3 replicates, 3 controls) were used in our analysis. For validation we used two additional datasets – Tyanova *et al* (Tyanova et al., 2016) containing 40 tumor proteomes from diverse breast cancer subtypes, and Pozniak *et al* (Pozniak et al., 2016) containing 66 proteomes from primary luminal-type breast tumors or metastases. The dataset of Tyanova *et al* contains SILAC ratios which we converted using a log2 transformation prior to calculating correlations. For all proteomics datasets proteins absent in more than 40% of samples were discarded. Multiple proteins mapping to the same gene were averaged into a single gene-level score. The resulting datasets contained profiles for 9,833 proteins (Mertins et al., 2016), 5,248 proteins (Tyanova et al., 2016) and 4,361 proteins (Pozniak et al., 2016).

### Protein Interaction Network

We assembled an integrated protein interaction network from multiple sources. From the HINT database (Das and Yu, 2012) we included all co-complex interactions that were reported in at least two publications. From the BioGRID database (Chatr-Aryamontri et al., 2016) we included all protein-protein interactions in the multi-validated interactome – a network of interactions that were either observed in two experimental systems or in two separate publications. We augmented this set of high-confidence interactions with the result of four recent large-scale protein interactome mapping efforts (Havugimana et al., 2012; Hein et al., 2015; Huttlin et al., 2015; Wan et al., 2015). The resulting integrated network contained 83,656 interactions between 11,930 proteins.

### Identifying Protein Complexes

Our goal was to identify sets of proteins (complexes) such that each complex consisted of a set of proteins whose expression profiles were highly similar across tumor profiles and that were densely connected on the protein interaction network. Other formulations are possible, but we chose to focus on disjoint complexes, such that each protein could only belong to a single complex. We did not require that every protein be assigned to a complex.

There are three components to our approach 1) choosing a score to evaluate the similarity of the expression profiles of a set of proteins 2) the identification of a similar score to evaluate the connectivity of a set of proteins on an interaction network, and 3) the identification of sets of proteins that score well on both datasets.

#### 1) Scoring complexes using expression profiles

We calculate the Pearson’s correlation coefficient between each pair of expression profiles (A, B) and use this to compute a log-likelihood ratio that A and B belong to the same protein complex versus the likelihood that they are unrelated. This can formalized as follows:

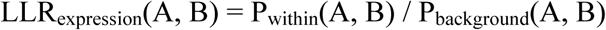

P_within_ is calculated using logistic regression trained on CORUM co-complexed pairs (Ruepp et al., 2010) as true positive examples. To prevent bias resulting from the large number of co-complex pairs falling within extraordinarily large complexes (e.g. Spliceosome, Proteasome, Ribosome) we exclude CORUM complexes containing more than 30 proteins from our training set. We assume a ratio of 300 negatives for every true positive, consistent with estimates of the size of the human interactome (Stumpf et al., 2008). Negative training examples are chosen randomly from the set of proteins with measured protein expression. P_background_ is the probability of observing the measured correlation between A and B in the set of all pairwise correlations.

For each set of proteins (S) we calculate the total LL_Rexpression_(S) as the sum of all LL_Rexpression_(A,B) scores for all unordered pairs (A,B) in the set S.

#### 2) Scoring complexes using the protein-protein interaction network

For the protein-protein interaction network we sought to score each pair of proteins based on how likely they are to form part of the same complex. While direct protein-protein interaction provides an indication that two proteins may be part of a protein complex, previous work has demonstrated that taking into account the fraction of interaction partners shared by two proteins can provide additional support of co-complex membership (Bader et al., 2004; Goldberg and Roth, 2003). Based on this principle we assigned a weighted score to every pair of interacting proteins in our integrated network accounting for the proportion of interaction partners they share. This score was equal to a –log10 transformed p-value calculated from a hypergeometric test that assessed the significance of the number of interaction partners they shared. An advantage of this approach is that two proteins that interact with each other directly and share all of their interaction partners will be given a higher score than two proteins that interact with each other but have no other interaction partners in common.

As with the protein expression correlation, this score was transformed into log-likelihood ratio (LLRinteraction) by comparing the probability of observing a particular score within a protein complex to the probability of observing it among all pairs of proteins. For each set of proteins (S) we calculate the total LLRinteraction(S) as the sum of all LLRinteraction(A,B) scores for all unordered pairs (A,B) in the set S.

#### 3) Identifying complexes supported by both data sources

For each set of proteins we can assign a score LLR_integted_(S), which is equal to the sum of LLR_expression_(S) and LLR_interaction_(S). Our challenge is the identification of sets of proteins with high LLR_integrated_ scores. As we are only interested in sets of proteins that score well on both resources we can restrict our search to those sets that have a positive LLR_interaction_ and a positive LLR_expression_ (i.e. we are only interested in sets of proteins that have highly correlated protein expression **and** are densely connected on the protein interaction network, not one or the other).

We identify high-scoring sets of proteins using an approach resembling agglomerative hierarchical clustering. Similar approaches have been used previously to identify complexes supported by genetic interaction and protein interaction networks in budding yeast (Bandyopadhyay et al., 2008) and also to identify complexes supported by the genetic interaction networks of two distinct yeast species (Ryan et al., 2012).

To initialize our clusters we first evaluate LLR_integrated_ for all pairs of proteins that directly interact in the protein-protein interaction network. We also evaluate scores for all possible 3-cliques (sets of three proteins that all interact with each other) in the protein-protein interaction network. The highest scoring pair or 3-clique is taken as an initial cluster, and all overlapping pairs or 3-cliques are then removed from consideration. The second highest scoring pair or 3-clique is then assigned as a cluster, and any overlapping pairs or 3-cliques removed from consideration. This continues until no pairs or 3-cliques with positive LLR_integrated_ scores remain. At the end of the process proteins that have not been assigned to any cluster are assigned to their own single element cluster. We then apply an iterative approach to improve these clusters. At each iteration we consider three possible moves – merging, removal and switching. Each pair of clusters (m_1_,m_2_) is evaluated for merging into a single cluster (m_1_ U m_2_) and assigned a score LLR_integrated_(m_1_,m_2_). For every protein in every cluster with multiple proteins we also calculate a LLR_remove_ score that reflects the change in the log likelihood resulting from removing that protein from the cluster, and an LLR_switch_ score that calculates the change in likelihood from switching a protein from one cluster to another. At each iteration max(LLR_merge_, LLR_remove_, LLR_switch_) is taken as the next move. To prevent the identification of clusters supported by only one data source (e.g. highly correlated expression but not densely connected on the protein interaction network) we only permitted moves in cases where the move resulted in an increase in the LLR score for both the expression and the protein interaction networks. Iterations continue until no move that increases the LLR score on both sources is identified. The end result is a list of clusters with an associated LLR score.

#### Estimating a protein complex false discovery rate

We assume that by chance some proteins that interact on the protein interaction network would have high co-expression scores and consequently we could identify clusters with positive LLR_expression_ and LLR_interaction_ scores. To remove potentially spuriously detected clusters we compared the clusters we identified to those identified using 10 randomized versions of the input - the same protein interaction network and expression set, but with the gene IDs on the expression set shuffled. These randomized networks allowed us to empirically estimate the False Discovery Rate as we could see for a given LLR_integrated_ score how many complexes would be discovered in the randomized networks. We chose an FDR of 10% for defining the BrCa-Core set of complexes.

### Evaluating Complexes

To assess the overlap between BrCa-Core complexes and existing annotation sets (CORUM complexes, Gene Ontology Cellular Compartment, Gene Ontology Biological Process) we used the gProfiler tool (Reimand et al., 2016). Only genes present in both the protein-interaction network and the tumor proteome expression were used as the background list or this enrichment. Multiple testing correction was performed using the default g:SCS approach (Reimand et al., 2016).

We calculated the average Pearson correlation between complex subunits using the dataset of Tyanova *et al* (Tyanova et al., 2016) and Pozniak *et al* (Pozniak et al., 2016). For this analysis we excluded pairs of proteins whose genes reside on the same chromosome to avoid high correlation resulting solely from co-amplification/codeletion events. For the shRNA data from (Marcotte et al., 2016) we calculated the Pearson’s correlation of co-complexed pairs using the zGARP profiles of 77 breast cancer cell lines.

From the STRING database (Szklarczyk et al., 2017) we extracted pairs of proteins that are frequently mentioned together in the literature (textmining score > 250) and that tend to co-occur in a significant pattern across species (cooccurence score > 0). Fisher’s exact test was used to assess the significance of the overlap between the BrCa-Core co-complexed pairs and these reference datasets.

### Identifying subtype specific expression patterns

To identify protein complexes differentially expressed in specific breast cancer subtypes we used a variant of the 1D annotation enrichment test proposed by Cox and Mann (Cox and Mann, 2012). For each protein we calculate the difference between the median expression of samples from a specific subtype and the median expression of samples from all other subtypes combined. We then applied a Mann-Whitney test to these median differences to see if the members of a given protein complex are among the most significantly differentially expressed proteins in a particular subtype (i.e. to see if all/most complex members are at one end of a ranked list of differentially expressed proteins). This test is performed in a two-sided fashion to identify complexes that are either over- or under-expressed in specific subtypes. All protein complexes with more than two members are tested for differential expression in all three subtypes. We correct for multiple-hypothesis testing using the Benjamini and Hochberg approach (Benjamini and Hochberg, 1995), and identified a set of 82 differentially expressed complexes at an FDR of 10%. We then tested these complexes for differential expression in the dataset of Tyanova *et al* at the same FDR. As not every BrCa-Core complex is represented by multiple members in Tyanova *et al* we could test only 59 of these associations. The s-score (Cox and Mann, 2012) was used to measure the effect size of the association between protein complex expression and subtype, and Spearman’s correlation was used to assess the concordance of effect sizes between the associations identified in the Mertins *et al* data and those in Tyanova *et al*.

### Mutation, copy number, mRNA expression and RPPA data

Sequence, copy number and mRNA expression profiles for were all obtained through the cBioPortal (Breast Invasive Carcinoma, TCGA Provisional) (Gao et al., 2013). To identify associations between mutation/deletion and protein abundance we annotated all tumor samples according to whether or not they featured mutations or deletions in each of the genes coding for proteins in the BrCa-Core set. For copy number profiles we considered genes to be deleted in a specific sample if they had a GISTIC score of - 2. We considered genes to be mutated if they harbored a non-synonymous missense mutation, splice-site mutation, an insertion or deletion, or a nonsense mutation. For the RPPA analysis and mRNA expression analysis presented in Figure 6 we used the Z-score normalized expression levels available through the cBioPortal (Gao et al., 2013).

### MCF7 Analysis

The MCF7 cell lines were grown in DMEM (Gibco) supplemented with 10% fetal bovine serum (Gibco) and 1% L-glutamine (Gibco).

#### Total lysate preparation for Mass spectrometry

Cells were plated in 100 mm dishes. Once confluent, media was discarded and cells were washed in PBS. Cells were lysed in a lysis buffer containing 2% SDS (Fisher Scientific), 20 mM Tris-HCl pH 7.5, 150 mM NaCl, 1 mM MgCl_2_, (Sigma Aldrich) supplemented with protease inhibitor tablets (Roche) and phosphatase inhibitors (2 mM sodium orthovanadate, 10 mM sodium fluoride and 10 mM *β* - glycerophosphate) (Sigma-Aldrich). Lysates were subjected to sonication (Syclon ultrasonic cell disrupter), boiling (95**°**C, 5 min) and placed on ice for 10-15 min prior to centrifugation (14000 rcf, 10 min). The supernatant was transferred to fresh eppendorfs and samples were subsequently placed on ice for a further 10-15 min to allow the SDS to precipitate and re-centrifuged. Supernatant was transferred to fresh eppendorfs and protein concentration was measured using the Pierce BCA protein assay kit as per manufacturers instruction (Thermo Scientific), using a SpectraMax M3 (Molecular Devices). Once quantified, DL-dithiothreitol (DTT) was added to the lysates at a final concentration of 0.1 M DTT. Subsequently, lysates were boiled (95**°**C, 5 min). Detergent was removed from the lysates prior to MS analysis using the Filter Aided Sample Preparation (FASP) procedure incorporating Vivacon spin ultracentrifugation units with a molecular weight cutoff of 30 kDa (Sartorius)(Wisniewski et al., 2009). Briefly, 200 *μ* l of urea buffer (Fisher Scientific) UA buffer (8 M urea in 0.1 M Tris-HCl pH 8.9) was added to 100 *μ* g of cell lysate. Samples were added to the filter unit and centrifuged at 14000 rcf for 15 min. An additional 200 *μ* l of UA buffer was added to the filter unit and re-centrifuged. Iodoacetamide (100 *μ* l, 0.05 M prepared in UA buffer) was added to the filter units, incubated for 1 min on a thermomixer at 600 rpm and subsequently incubated in darkness for 20 min. Following the incubation period, filter units were centrifuged and washed twice with 100 *μ* l of UA buffer followed by 2 washes with 100 *μ* l of ABC solution (0.05 M NH_4_HC0_3_). After the final wash step, filter units were transferred to a new collection tube and a multi-step digestion method was employed as described by Wisniewski and Mann (Wisniewski and Mann, 2012). In the first instance, proteins were digested in a wet chamber overnight at 37°C using a solution containing Lys-C (Lysl Endopeptidase, Wako) and ABC buffer (1:50, enzyme to protein ratio). The following day, liberated peptides were collected by centrifugation and subsequent wash cycles with ABC buffer. Meanwhile, remaining proteins on the filter unit were digested using a solution containing Sequencing Grade Modified Trypsin (Promega) and ABC buffer in a wet chamber at 37°C for a minimum of 4 hr. Once again liberated peptides were collected by centrifugation and subsequent wash cycles with ABC buffer. The concentration of the Lys-C digests and Trypsin digests were measured using a NanoDrop 2000. In total, 10 *μ* g of each digest was loaded onto activated handmade C18 StageTips as described previously(Rappsilber et al., 2003). StageTips were desalted with two 1% TFA wash cycles and bound peptides were eluted with 2 X 25 *μ* l of 50% ACN/0.1 % TFA. Final eluates were concentrated in the speed-vacuum centrifuge (Centri-Vap concentrator, Labconco to a final volume of ~5 *μ* l. Samples were then resuspended by adding 0.1% acetic acid, to a final volume of 15 *μ* l and analyzed by mass spectrometry.

#### Mass Spectrometry

Mass spectrometry analysis was performed on a Q-Exactive mass spectrometer (Thermo Scientific), connected to a Dionex Ultimate 3000 (RSLCnano) chromatography system (Thermo Scientific) incorporating an autosampler. Five microliters of Lys-C/tryptic peptides was loaded onto a fused silica emitter (75 *μ* m ID, pulled using a laser puller (Sutter Instruments P2000)), packed with 1.8 *μ* 120Å UChrom C18 packing material (NanoLCMS Solutions) and separated using an increasing acetonitrile gradient of 2 – 35%, with a 180 min reverse phase gradient at a flow rate of 250 nl/min. The instrument was operating in positive ion mode and with a capillary temperature of 320**°**C, coupled to a potential of 2300V applied to the column. Scan parameters for MS1 were as follows:Resolution 70,000, AGC 3e^6,^ MIT 60ms while scan parameters for MS2 were: Resolution 17,500, AGC 5e^4^, MIT 250ms, NCE 27.0, Isolation window 1.6m/z. The exclusion list parameters contained no entries and charge exclusion was set to un-assigned and singly charged. Both MS1 and MS2 were recorded as profile data. Data were acquired in automatic data-dependent switching mode, with a high-resolution MS scan (300-1600 m/z) selecting the 12 most intense ions prior to tandem MS (MS/MS) analysis. Each biological sample (n=3) was run in technical duplicate. The resulting mass spectra were analyzed using MaxQuant software (version 1.5.0.25) containing the in-built Andromeda search engine to identify the proteins from a human database (Uniprot HUMAN, release 2012_01) containing 20,242 entries. Default parameters were selected in MaxQuant with the exception of the selection of the relevant enzyme, (LysC and Trypsin digests were separated between parameter groups). For database searches, the precursor mass tolerance was set to 20 ppm for first searches and 4.5 ppm for main Andromeda search. The search included a fixed modification of Carbamidomethyl (C) and variable modifications of Oxidation (M);Acetyl (Protein N-term). Label free quantification with a minimum ratio count of 2 was selected, the maximum number of missed cleavages was set at 2 and minimum peptide length was set to 7 amino acids. An FDR of 0.01 was set for peptide and protein identifications. Match between runs was selected with a matching time window of 0.7 min and alignment time window of 20min. The presence of reverse and contaminant identifications were removed from the dataset.

#### Differential expression analysis

Proteomic profiles were generated for three biological replicates of the parental (*CDH1* wild-type) and *CDH1*-defective cell lines. Two technical replicates were obtained for each biological replicate and these were averaged prior to further analysis. Missing values were imputed using the minimum observed intensity for each sample, based on the assumption that missing proteins could be absent or below the detection threshold of the instrument. Log2 transformed LFQ (Label Free Quantification) values were used for analysis. A two-sided heteroscedastic t-test (Welch’s t-test) was used to identify differentially expressed proteins and the Benjamini-Hochberg approach was used to estimate the False Discovery Rate (Benjamini and Hochberg, 1995).

## Acknowledgements

We thank Denis Shields, Ariane Watson and Gerard Cagney for comments on the manuscript and Pedro Beltrao for helpful discussion. Data used in this publication were generated by the Clinical Proteomic Tumor Analysis Consortium (NCI/NIH). CJR is a Sir Henry Wellcome Fellow jointly funded by Science Foundation Ireland, the Health Research Board, and the Wellcome Trust (grant number 103049/Z/13/Z) under the SFI-HRB-Wellcome Trust Biomedical Research Partnership. CJL is supported by Cancer Research UK (grant number C347/A8363) and Breast Cancer Now. We acknowledge NHS funding to the ICR/Royal Marsden Hospital Biomedical Research Centre.

## Supplemental Figure Legends

**Figure S1.**
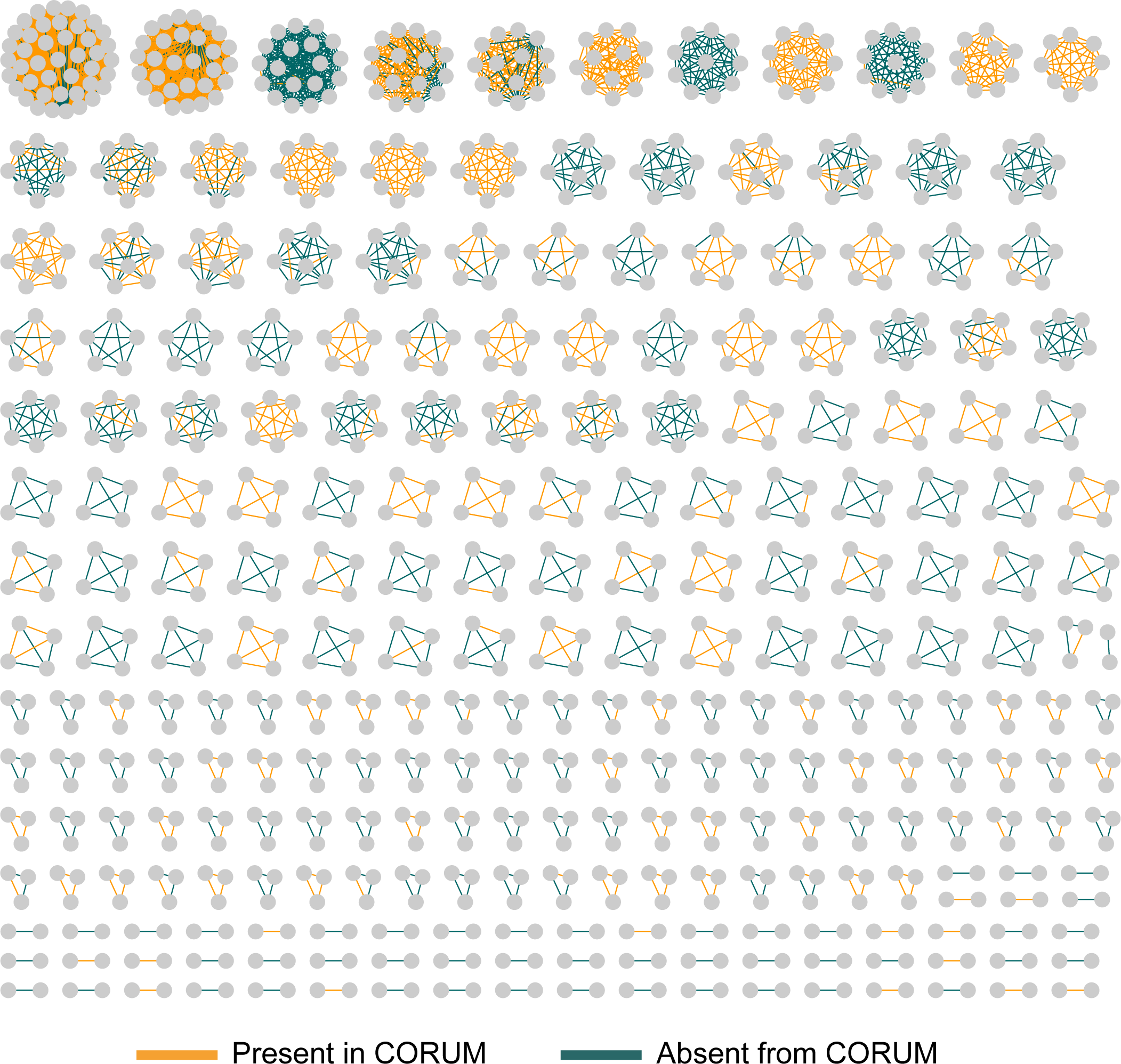
BrCa-Core Complexes. Related to Figure 3. All complexes identified in BrCa-Core are shown. Orange edges correspond to co-complex relationships also identified in the CORUM database while green edges correspond to co-complex relationships absent from CORUM.

**Figure S2.**
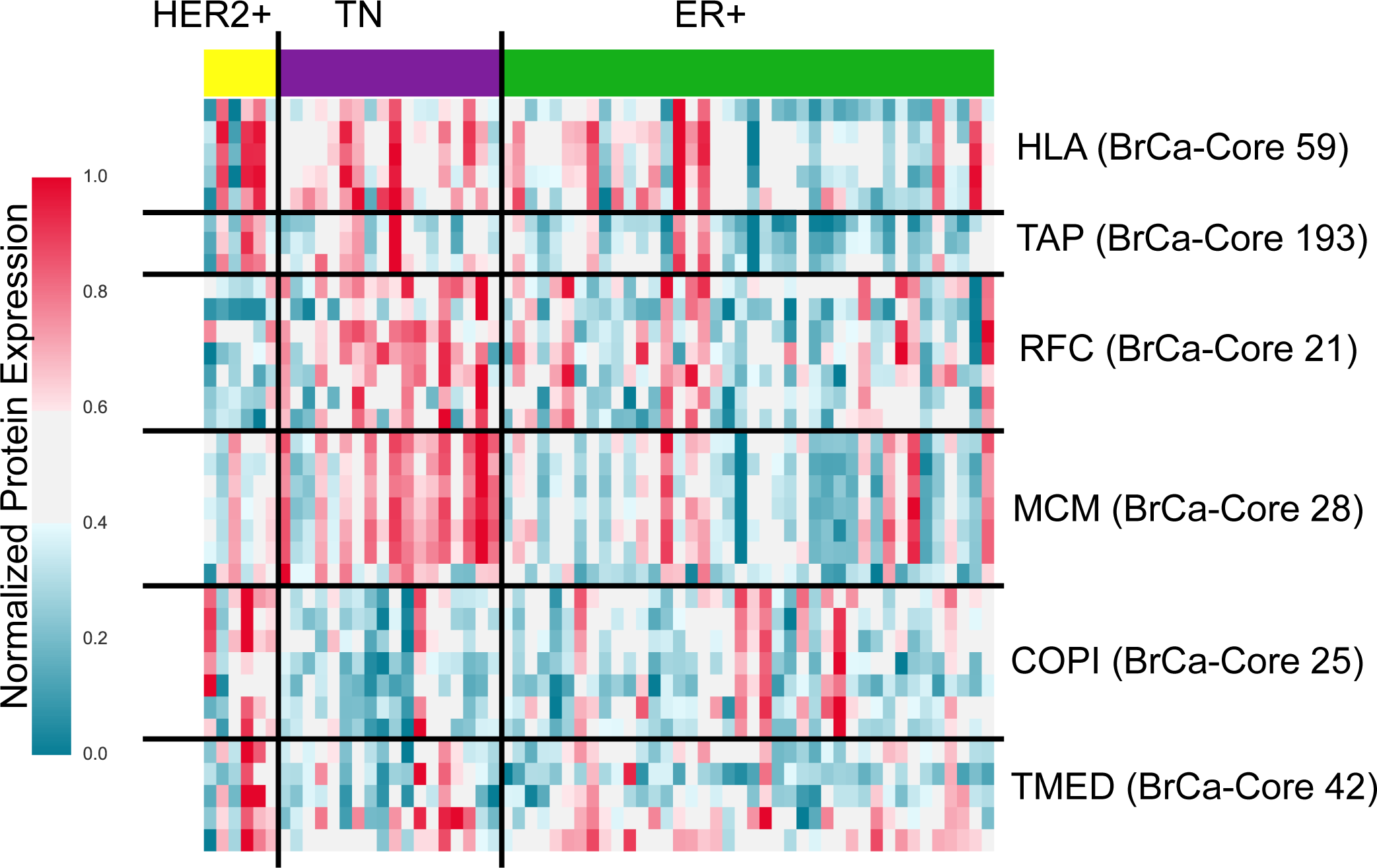
Subtype specific complex expression. Related to Figure 4. Heatmap displaying expression levels of specific protein complexes. Tumor samples are grouped according to subtype (using IHC), indicated on top of the heatmap. Genes are grouped into specific complexes indicated on the right of the heatmap. All expression levels have been normalized such that the maximum expression level is 1 and minimum is 0. Shown are the expression levels from Mertens *et al* (used for discovery). Expression levels for Tyanova *et al* (used for validation) are shown in Figure 4

**Figure S3.**
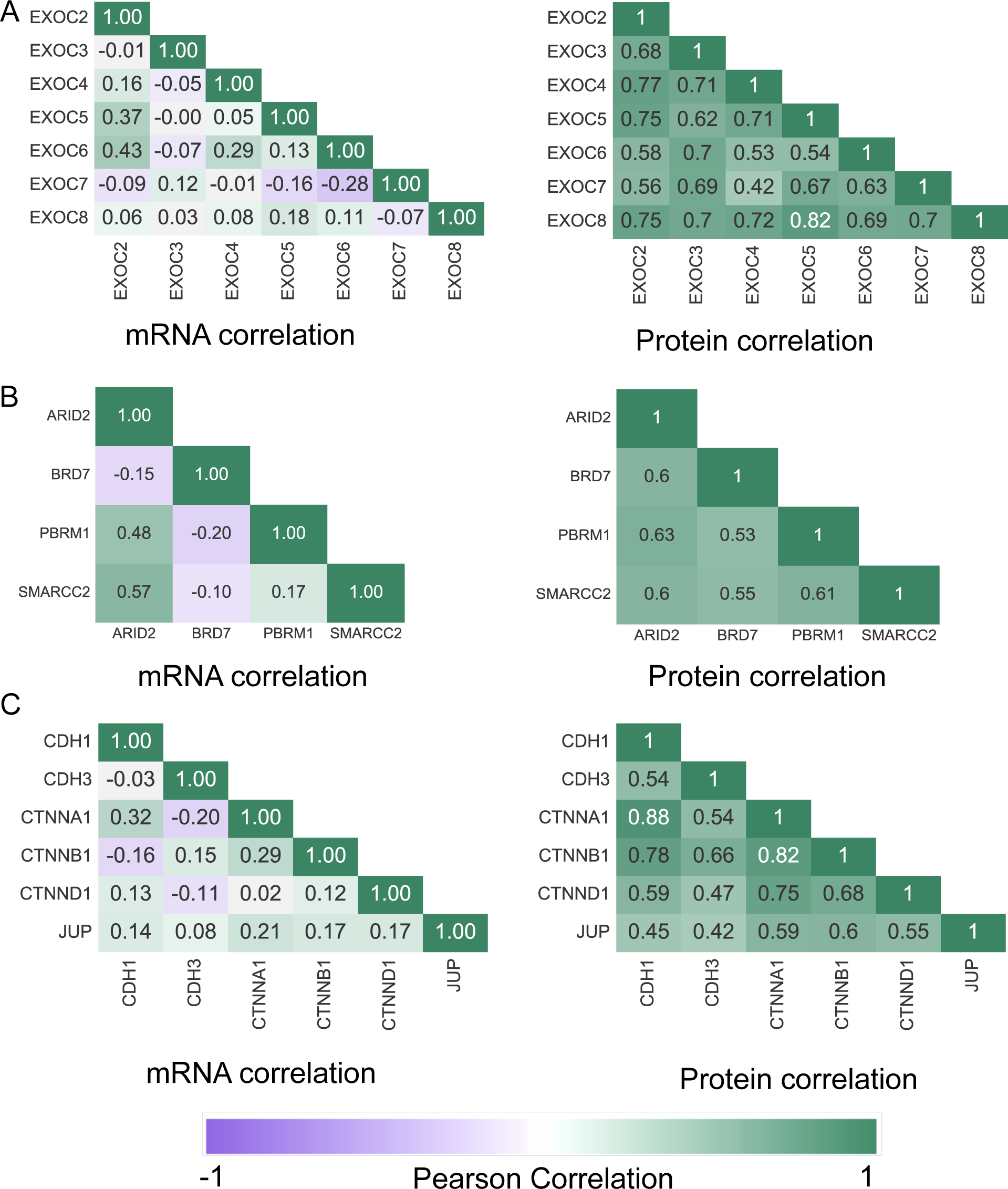
Correlation within BrCa-Core complexes is more evident using proteomic profiles than mRNA profiles. Related to Figure 5. Heatmaps showing the Pearson’s correlation between the subunits of the (A) exocyst subcomplex (main text Figure 5B), (B) BAF complex (main text Figure 5C) and (C) the adherens junction complex (main text Figure 5D). In each subfigure the heatmap calculated using mRNA profiles is on the left and using protein profiles on the right

**Figure S4.**
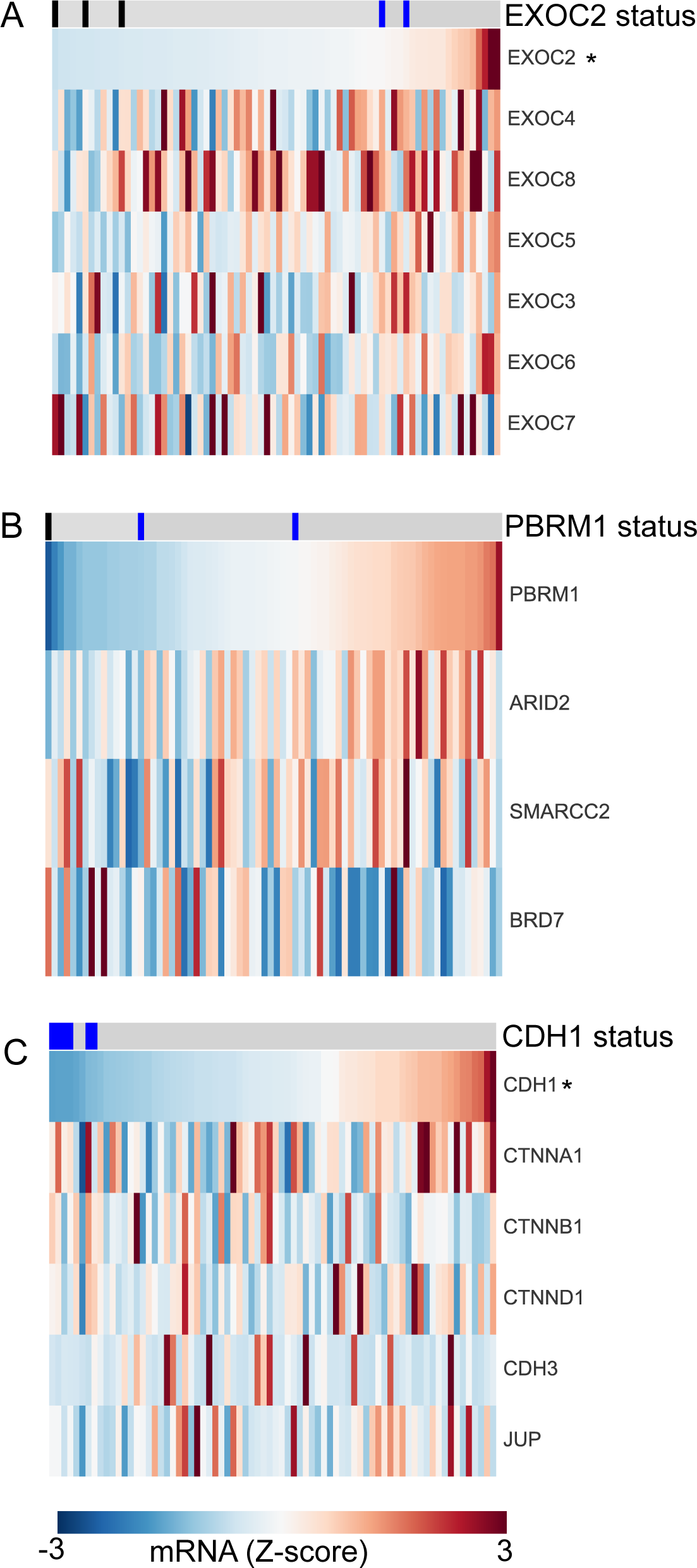
Subunit loss is not associated with a reduction in protein complex mRNA expression. Related to Figure 5. A) Mutation or deletion of *EXOC2* is not associated with a reduction in mRNA expression of the exocyst complex. *EXOC2* mutation (blue) or deletion (black) is indicated in the top row. Samples have been sorted according to EXOC2 mRNA expression. EXOC2 (marked with a star) is the only gene to display a significant association with *EXOC2* status (P<0.05, Mann-Whitney U test) B) Mutation or deletion of *PBRM1* is not associated with a reduction in mRNA expression of the BAF subcomplex members. No member of the complex displays a significant association between mRNA and *PBRM1* status. Legend as for A. C) *CDH1* mutation is not associated with a reduction in mRNA expression of an adherens-junction complex. Legend as for A. CDH1 (marked with a star) is the only gene to display a significant association with *CDH1* status (P<0.05, Mann-Whitney U test).

